## Supplementary file 1 for "A simple mechanism for integration of quorum sensing and cAMP signalling in *V. cholerae*"

**Table S1: HapR binding targets identified by previous studies**

| gene(s) | evidence[1] | source | detected by ChIP-seq[2] |
|---|---|---|---|
| *VC0583* | EMSA, DNAse I footprint | (1) | yes |
| *VC2647* | EMSA, DNAse I footprint | (2) | no |
| *VCA0865* | not shown directly | (3) | yes (below cut-off) |
| *VCA0952* | EMSA | (4) | yes (below cut-off) |
| *VC2370* | EMSA | (4) | no |
| *VC0900* | EMSA | (4) | no |
| *VC1851* | EMSA | (4) | yes |
| *VC1086* | EMSA | (4) | no |
| *VCA0074* | EMSA | (4) | no |
| *VCA0246<>VCA0247* | EMSA | (5) | yes (below cut-off) |
| *VC1222* | EMSA | (5) | no |
| *VC1181* | EMSA | (5) | no |
| *VC0432* | EMSA | (5) | no |
| *VC2762* | EMSA | (5) | no |
| *VC1000* | EMSA | (5) | no |
| *VC2634<>VC2635* | EMSA | (5) | no |
| *VCA0017* | EMSA | (5) | yes (below cut-off) |
| *VCA0182<>VCA0183* | EMSA | (5) | no |
| *VC0166<>VC0167* | EMSA | (5) | no |
| *VC1415* | EMSA | (5) | yes (below cut-off) |
| *VCA0880* | EMSA | (5) | yes (below cut-off) |
| *VCA0684* | EMSA | (5) | no |
| *VC2674* | EMSA | (5) | no |
| *VC1213* | EMSA | (5) | yes (below cut-off) |
| *VC2035* | EMSA | (5) | no |
| *VC0089* | EMSA | (5) | no |
| *VCA0865* | EMSA | (5) | yes (below cut-off) |
| *VCA0148* | EMSA | (5) | yes |
| *VC0934* | EMSA | (5) | no |
| *VC0241* | EMSA | (5) | yes |

[1]The column lists evidence for direct binding either using electrophoretic mobility shift assay (EMSA) or DNAse I footprinting.

[2]The column lists evidence for binding from ChIP-seq assays. In cases where a binding peak was evident upon visual inspection, but this fell below our stringent cut-off, this is indicated.
