## Supplementary file 2 for "A simple mechanism for integration of quorum sensing and cAMP signalling in *V. cholerae*"

**Table S2:** Strains, plasmids, and oligonucleotides

| Name | Description | Source |
|---|---|---|
| | **Bacterial Strains** | |
| | *Vibrio cholerae*[1] | |
| E7946 | Wildtype *Vibrio cholerae* El Tor Ogawa derivative, SmR | (1) |
| E7946 Δ*hapR* | E7946 derivative lacking *hapR* | This work |
| E7946 Δ*crp* | E7946 derivative lacking *crp* | This work |
| E7946 Δ*hapR*Δ*crp* | E7946 derivative lacking *hapR* and *crp* | This work |
| E7946 Δ*luxO* | E7946 derivative lacking *luxO* | This work |
| E7946 Δ*hapR* | E7946 derivative *hapR::Spec*$^{R}$ | This work |
| E7946 Δ*luxO*Δ*hapR* | E7946 derivative lacking *luxO*, *hapR::Spec*$^{R}$ | This work |
| E7946 Δ*murP* | E7946 derivative *lacking murP* | (2) |
| | *Escherichia coli* | |
| DH5α | *fhuA*2 Δ(*argF*-*lacZ*)U169 *phoA glnV*44 Φ80 Δ(*lacZ*)M15 *gyrA*96 *recA*1 *relA*1 *endA*1 *thi*-1 *hsdR*17 | NEB |
| T7 Express | *fhuA2 lacZ::T7 gene1 [lon] ompT gal sulA11 R(mcr-73::mini Tn10*--Tet$^{S}$*)2 [dcm] R(zgb-210::Tn10*--Tet$^{S}$*) endA1 Δ(mcrC-mrr)114::IS10* | NEB |
| JCB387 | Δ*nirB*, Δ*lac* | (3) |
| S17 λpir | *lac*U169 (*lac*ZM15), *recA*1, *endA*1, *hsdR*17, *thi*-1, *gyrA*96, *relA*1, *pir* | (4) |
| | **Plasmids** | |
| pSR | pBR322-derived plasmid. Features cloning site upstream λoop transcription terminator. AmpR | (5) |
| pRW50T | A broad-host range *lacZ* expression vector encoding *oriT* of pRK, TetR, Tra+ | (6) |
| pAMNF | Plasmid for basal expression of N-terminal 3xFLAG-tagged proteins, KanR | (7) |
| pAMCF | Plasmid for basal expression of C-terminal 3xFLAG-tagged proteins, KanR | (7) |
| pRK2013 | Helper plasmid for conjugation, KanR, oriColE1, RK2-, Mob+, Tra+ | (8) |
| pKAS32 | Suicide plasmid for mutant strain construction, AmpR | (9) |
| | **Oligonucleotides**[2,3] | |
| | Oligonucleotides for cloning *hapR* or *luxO* (5' to 3') | |
| *hapR*\|pAMNF\|fwd | GGCTGC*GGTACC*ATGGACGCATCAATCGAAAAACGC | This work |
| *hapR*\|pAMNF\|rev | GCCCG*AAGCTT*CTAGTTCTTATAGATACACAGCATATTGAGG | This work |

| | | |
|---|---|---|
| *luxO*\|pAMCF\|fwd | GGCTGC*GGTACC*ATGGTAGAAGACACGGCGTCGGTG GCGGCGCTGTATCGTTCTTACCTCACACCGCTGGATA TTGATATCAATATCGTGGGTACGGGAC | This work |
| *luxO*\|pAMCF\|rev.1 | ATCGCGTCCCGTACCCACGATATTG | This work |
| *luxO*\|pAMCF\|rev.2 | GCCCG*AAGCTT*CCGTTCCTTCTCTTTTTCTTTCAC | This work |
| Oligonucleotides for cloning regulatory regions in pRW50T and pSR (5' to 3') | | |
| P*murQ*\|fwd | AAAA*GAATTC*CACCAATCTGGCGGCCACTC | This work |
| P*murQ*\|rev | TTTT*AAGCTT*CATAAGGCTTCTCGGCAAAT | This work |
| P*hapR*\|fwd | AAAA*GAATTC*CATACCATTCTCGTTGTGTT | This work |
| P*hapR*\|rev | TTTT*AAGCTT*CATAGGGGTATATCCTTGCC | This work |
| P*VC0585*\|fwd | AAAA*GAATTC*CATAGGGGTATATCCTTGCC | This work |
| P*VC0585*\|rev | TTTT*AAGCTT*CATACCATTCTCGTTGTGTT | This work |
| P*VC0502*\|fwd | AAAA*GAATTC*GACAAAAGTTTGTTGCCCGC | This work |
| P*VC0502*\|rev | TTTT*AAGCTT*CATAGTATTGGCTTTGGCAT | This work |
| P*mutH*\|fwd | AAAA*GAATTC*ACCAGACCACATGAAGATCC | This work |
| P*mutH*\|rev | TTTT*AAGCTT*CATAAAGGCTTTCGGTTTGG | This work |
| P*leuO*\|fwd | AAAA*GAATTC*TAATGAACTGACTAACTCA | This work |
| P*leuO*\|rev | TTTT*AAGCTT*CATTGCGTCTTTTTTATCT | This work |
| P*VC0433*\|fwd | AAAA*CAATTG*TACCTGCAACTTCAAGTAGT | This work |
| P*VC0433*\|rev | TTTT*AAGCTT*CATTATTTTTTAATCACCAA | This work |
| P*VC0241*\|fwd | AAAA*GAATTC*CTATAGATGCGAACAGTTGC | This work |
| P*VC0241*\|rev | TTTT*AAGCTT*CATAGTATGCTGACTACTGC | This work |
| P*rfaD*\|fwd | AAAA*CAATTG*CATAGTATGCTGACTACTGC | This work |
| P*rfaD*\|rev | TTTT*AAGCTTC*ATTATGAATTCCTATAGAT | This work |
| P*VC0620*\|fwd | AAAA*GAATTC*AAACCGTACCCGTTTTGCGAG | This work |
| P*VC0620*\|rev | TTTT*AAGCTT*ACTAGTAAGGAACAGCTATG | This work |
| P*VC0688*\|fwd | AAAA*GAATTC*CATAGCGTTTGTCCTTTTGT | This work |
| P*VC0688*\|rev | TTTT*AAGCTTC*ATTACTATCCATTTTTTCA | This work |
| P*VC1298*\|fwd | AAAA*GAATTC*CATAAACTCTTTGTATAATT | This work |
| P*VC1298*\|rev | TTTT*AAGCTT*CATGATAGTTTTGTAATTAT | This work |
| P*VC1375*\|fwd | AAAA*CAATTG*CATCCACTTCTTCCTTATTA | This work |
| P*VC1375*\|rev | TTTT*AAGCTT*CATATCAAAAGTGGTTGGGA | This work |
| P*VC1376*\|fwd | AAAA*CAATTG*CATATCAAAAGTGGTTGGGA | This work |
| P*VC1376*\|rev | TTTTAAGCTTCATCCACTTCTTCCTTATTA | This work |
| P*VC1403*\|fwd | AAAA*GAATTC*CATTTTTTGGGTAATCGATA | This work |
| P*VC1403*\|rev | TTTT*GGATCC*CATGGAAAACCTCGTTGTTT | This work |
| P*VC1405*\|fwd | AAAA*GAATTC*CATGGAAAACCTCGTTGTTT | This work |

| | | |
|---|---|---|
| P*VC1405*\|rev | TTTT*GGATCC*CATTTTTTGGGTAATCGATA | This work |
| P*VC1436*\|fwd | AAAA*GAATTC*CCAATTCGTAAAGAAGCTAA | This work |
| P*VC1436*\|rev | TTTT*GGATCC*CATATCTGCTGAGGCTTTAG | This work |
| P*VC2352*\|fwd | AAAA*GAATTC*TCAGCACAATATCTCGCGCC | This work |
| P*VC2352*\|rev | TTTT*AAGCTT*CATGCCGATGAGGCTCATAA | This work |
| P*VCA0218*\|fwd | AAAA*GAATTC*CATATAAACCTCACTGACTC | This work |
| P*VCA0218*\|rev | TTTT*AAGCTT*CATCGTTTACTTCTTATCAT | This work |
| P*VCA0219*\|fwd | AAAA*GAATTC*CATCGTTTACTTCTTATCAT | This work |
| P*VCA0219*\|rev | TTTT*AAGCTT*CATATAAACCTCACTGACTC | This work |
| P*VCA0662*\|fwd | AAAA*GAATTC*TCAGTTTGTAAATTGAGGAA | This work |
| P*VCA0662*\|rev | TTTT*AAGCTT*CATATCAAATATGAATCCTT | This work |
| P*VCA0663*\|fwd | AAAA*GAATTC*CATATCAAATATGAATCCTT | This work |
| P*VCA0663*\|rev | TTTT*AAGCTT*CATGGTGTTACCTACTTGTT | This work |
| P*VCA0906*\|fwd | AAAA*GAATTC*ATTTTTATCCTGACCCCATA | This work |
| P*VCA0906*\|rev | TTTT*GGATCC*CATGCCTACGCCTATCGCCG | This work |
| P*VCA0960*\|fwd | AAAA*GAATTC*CATCATGAGTTATATTTACA | This work |
| P*VCA0960*\|rev | TTTT*AAGCTT*CATCCCTCAATCCTCAGTTT | This work |
| P*VCA0961*\|fwd | AAAA*GAATTC*CATCCCTCAATCCTCAGTTT | This work |
| P*VCA0961*\|rev | TTTT*AAGCTT*CATCATGAGTTATATTTACA | This work |

Oligonucleotides for sequencing cloned DNA in plasmid constructs (5' to 3')

| | | |
|---|---|---|
| pRW50\|seq\|fwd | GTTCTCGCAAGGACGAGAATTTC | This work |
| pRW50\|seq\|rev | GTCGTTGAACTGAGCCTGAAATTCAGG | This work |
| pSR\|seq\|fwd | GTGCCACCTGACGTCTAAGAAACC | This work |
| pSR\|seq\|rev | GCAACCGAGCGTTCTGAACAAATCC | This work |
| pAM\|seq\|fwd | ATTTATTCCAATGTCACACACTTTTCGC | This work |
| pAM\|seq\|rev | GAAACGCCGTAGCGCCGATGGTAGT | This work |

Oligonucleotides for amplifying DNA for radio labelling (5' to 3')

| | | |
|---|---|---|
| pSR\|footprinting\|fwd | [BIOTIN]-CACGAGGCCCTTTCGTCTTCTC | This work |
| pSR\|footprinting\|rev | GGAGTTCTGAGGTCATTACTGGAG | This work |
| pSR\|footprinting\|rev | CACGAGGCCCTTTCGTCTTCTC | This work |
| pSR\|footprinting\|rev | [BIOTIN]-GGAGTTCTGAGGTCATTACTGGAG | This work |

Oligonucleotides for cloning DNA fragments in pKAS32 (5' to 3')

| | | |
|---|---|---|
| pKAS32\|fwd | GCAGGCACAAGCGGCCGCCTGCAGCTGGCGCCA TCGATACGCGTACGTCG | This work |
| pKAS32\|rev | CACGGTTTCATTAACAACCGGTACCTCTAGAACT | This work |

| | | |
|---|---|---|
| | ATAGCTAGCATGCGCAAATTTAAAGCGCTG | This work |
| *crp*\|arm\|up\|fwd | CGGTTGTTAATGAAACCGTGGATATTAAATGC | This work |
| *crp*\|arm\|up\|rev | ATCGGGGCACCTAGCCGATTTTTCCGGTTTC | This work |
| *crp*\|arm\|down\|fwd | AATCGGCTAGGTGCCCCGATAACCCGTC | This work |
| *crp*\|arm\|down\|rev | AGGCGGCCGCTTGTGCCTGCGCAGCCAA | This work |
| *hapR*\|arm1\|fwd | CTAGAGGTACCGGTTGTTAATCCCAACCCCGATT GGTAATC | This work |
| *hapR*\|arm1\|rev | TGTGTTTCATTTTCTTGGGCAGCACAAAG | This work |
| *hapR*\|arm2\|fwd | GCCCAAGAAAATGAAACACACAGTTGAAGTC | This work |
| *hapR*\|arm2\|rev | CGCCAGCTGCAGGCGGCCGCTATGCGGTCGATGT GCTGA | This work |
| | Oligonucleotides for mutagenesis of *crp* or *hapR* (5' to 3') | |
| hapR\|R123E\|fwd | TGCTTCAACCGAGGACGAAGTTTGGC | This work |
| hapR\|R123A\|fwd | TGCTTCAACCGCTGACGAAGTTTGGCC | This work |
| hapR\|R123\|rev | CTCCACTCAAACCAGACTTTG | This work |
| *crp*\|E55R\|fwd | GATCAAAGATCGTGAAGGTAAAGAGATGATTCTC | This work |
| *cpr*\|E55A\|fwd | GATCAAAGATGCGGAAGGTAAAGAGATGATTC | This work |
| *crp*\|E55\|rev | AGTACCGCAACTGAACCT | This work |

[1]E7946 Δ*luxO*, Δ*hapR*, Δ*lux*OΔ*hapR* and Δ*murP* derivatives are synonymous with strains SAD764, SAD793, TND3765 and SAD268.

[2]sequences in italic are sites for restriction endonucleases for use during cloning

[3]underlined sequences encode amino acid substitutions or remove unwanted sites for restriction endonucleases
