## Supplementary material for "A simple mechanism for integration of quorum sensing and cAMP signalling in *V. cholerae*": Figure 1-figure supplement 1

**a**

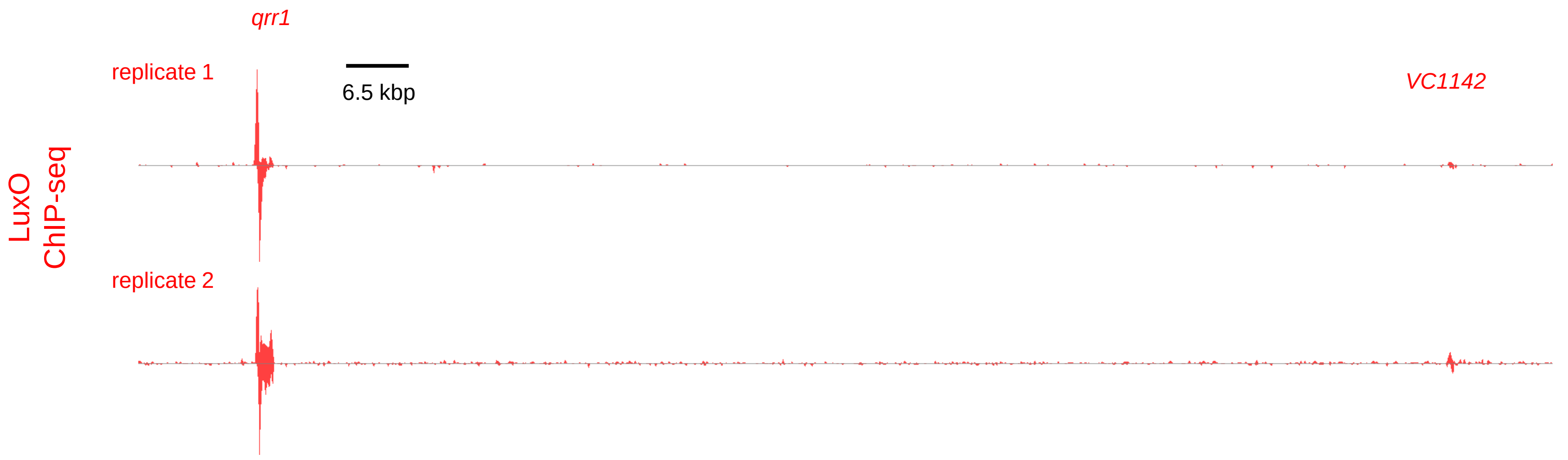


**b**

tata**cat**ccctc**A**tgcattttataactgatgttttaagagaaacgtcaatttgtccctaggacttcctgacttctcaaaagtgaata

gcgattcagcctatgcttaacgttagtcac**T**aattgacc**aattta**gacgatgattgaacgcagtcaataggacgaacgtataaaagc

acacagtttgatatgcaaactcaattttg**C**aattgca**attata**taaatttgtcat**ttgcaaatcgcga**agcaaatcattaacaagct

tgcatattggtgtcggtaaagtaagcaatcgcgctattcgtgatcgtcattcgcaatgttttatttgcagcgt**tacaat**tgctgc**A**t

**tagtct**ctaagt**A**agaggcgaaattcattaaagtgcggatgccgtcagtgagtttaaacggctagttttacagcactttatagttga

attctctgtaagaagccctcaaatattgaagt**G**tc**atg**
