## Supplementary figures and images for "A simple mechanism for integration of quorum sensing and cAMP signalling in *V. cholerae*"

### Figure 1-figure supplement 2

#### Figure 1-figure supplement 2

**a**

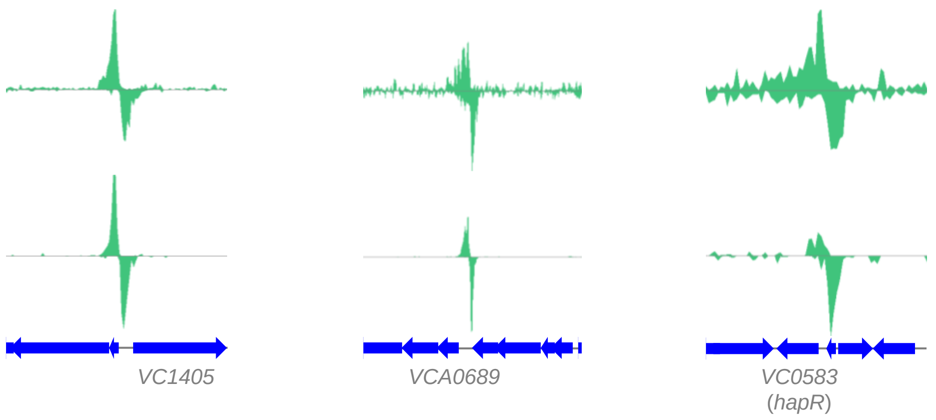


**b**

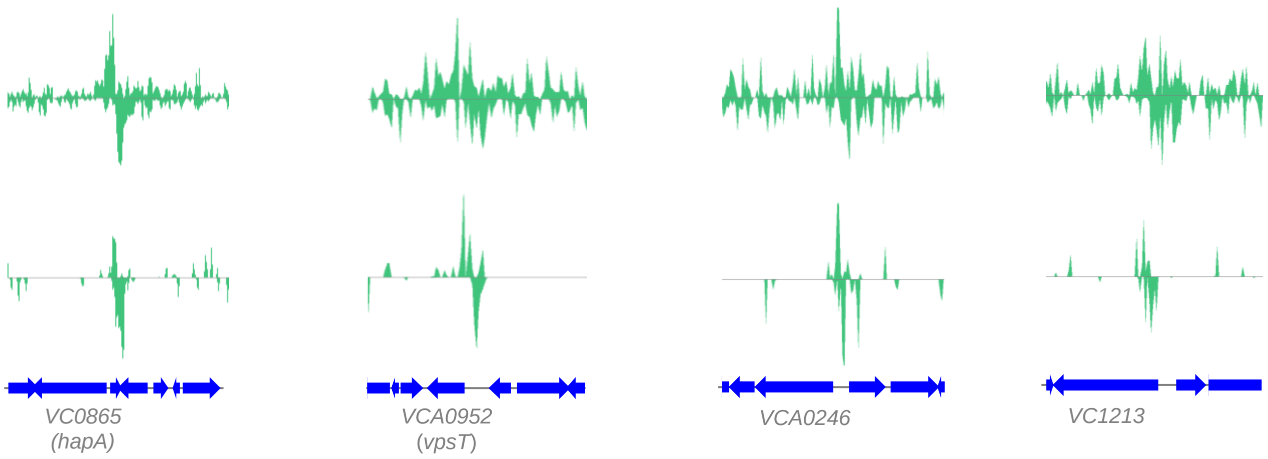

### Figure 2b source data 1

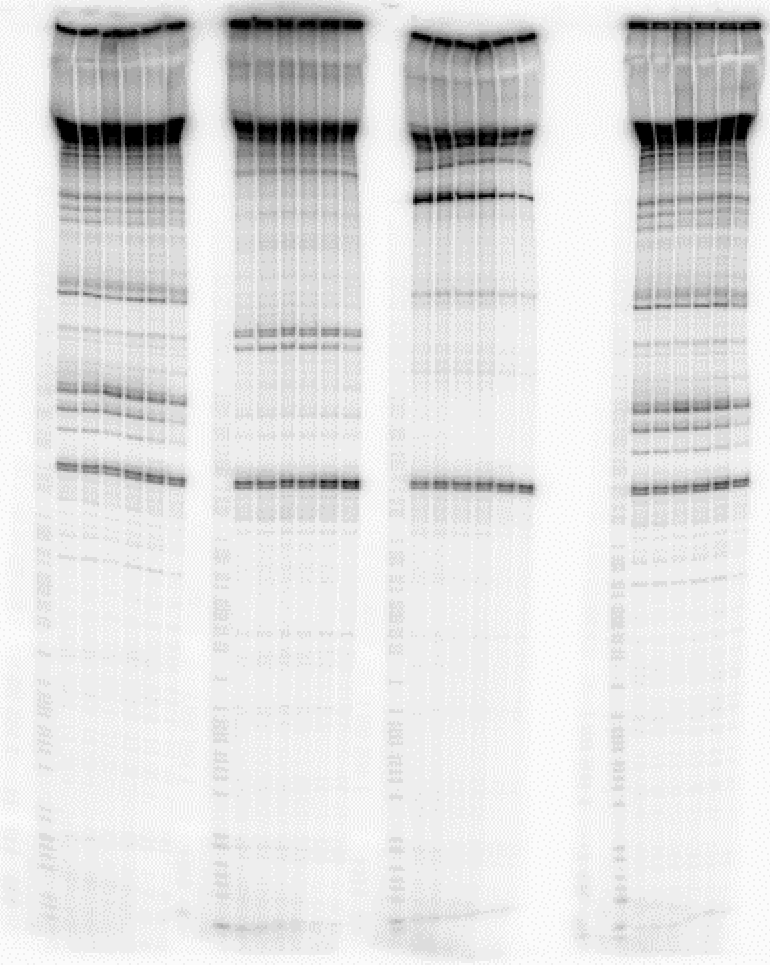

### Figure 2b source data 2

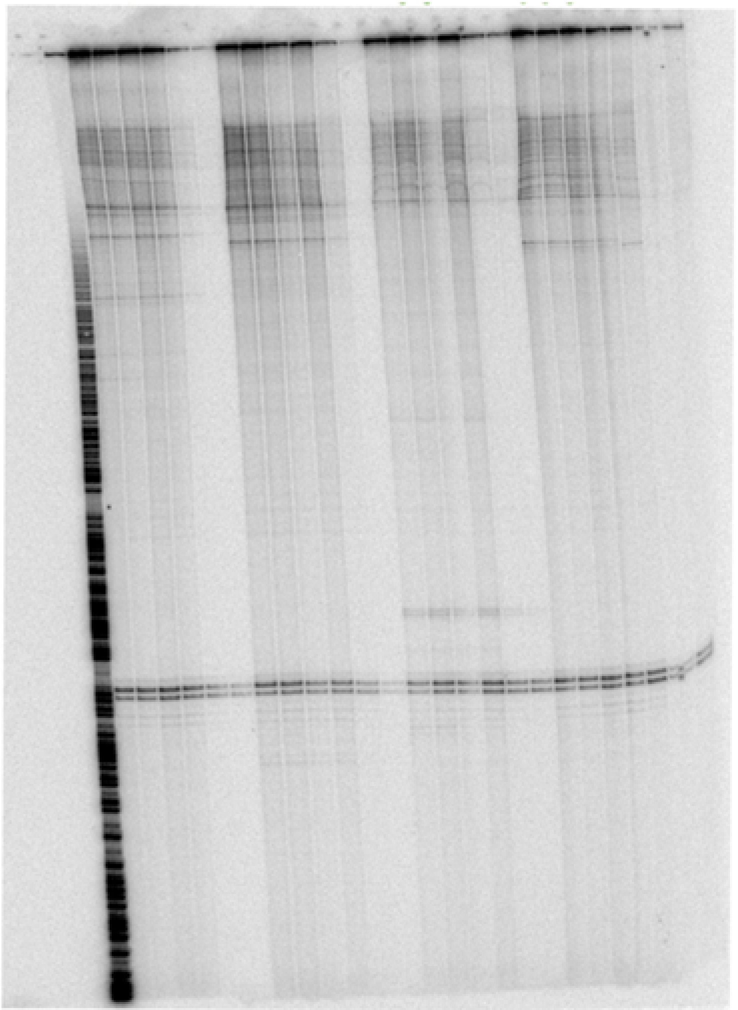

### Figure 2b source data 3

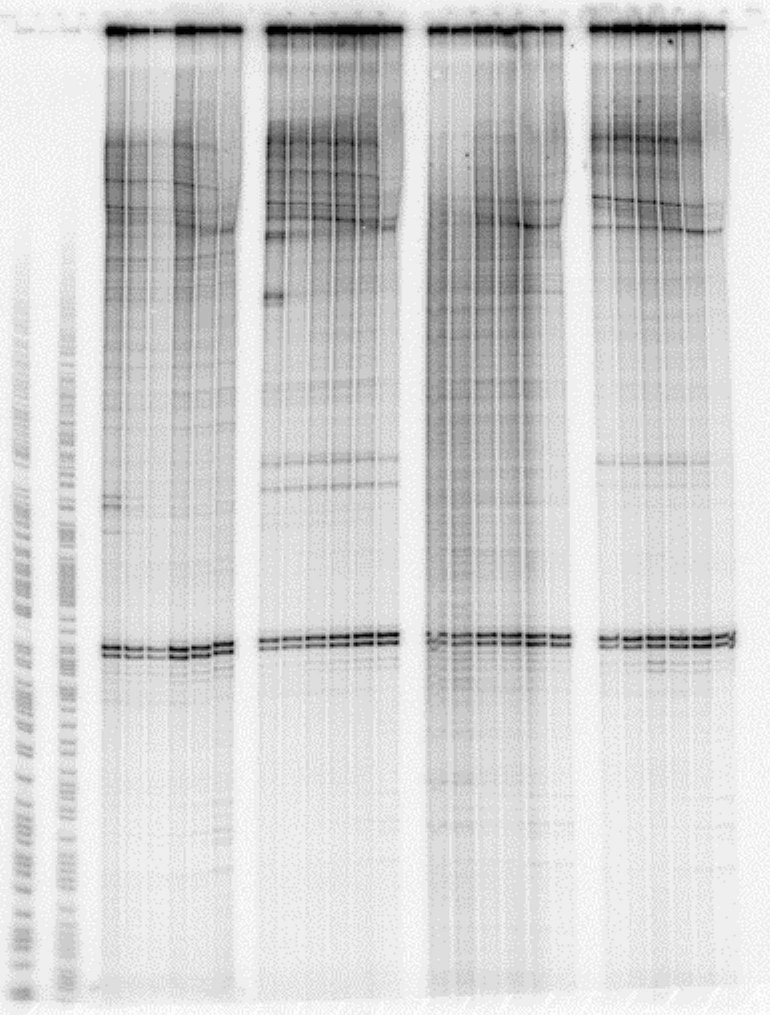

### Figure 2b source data 4

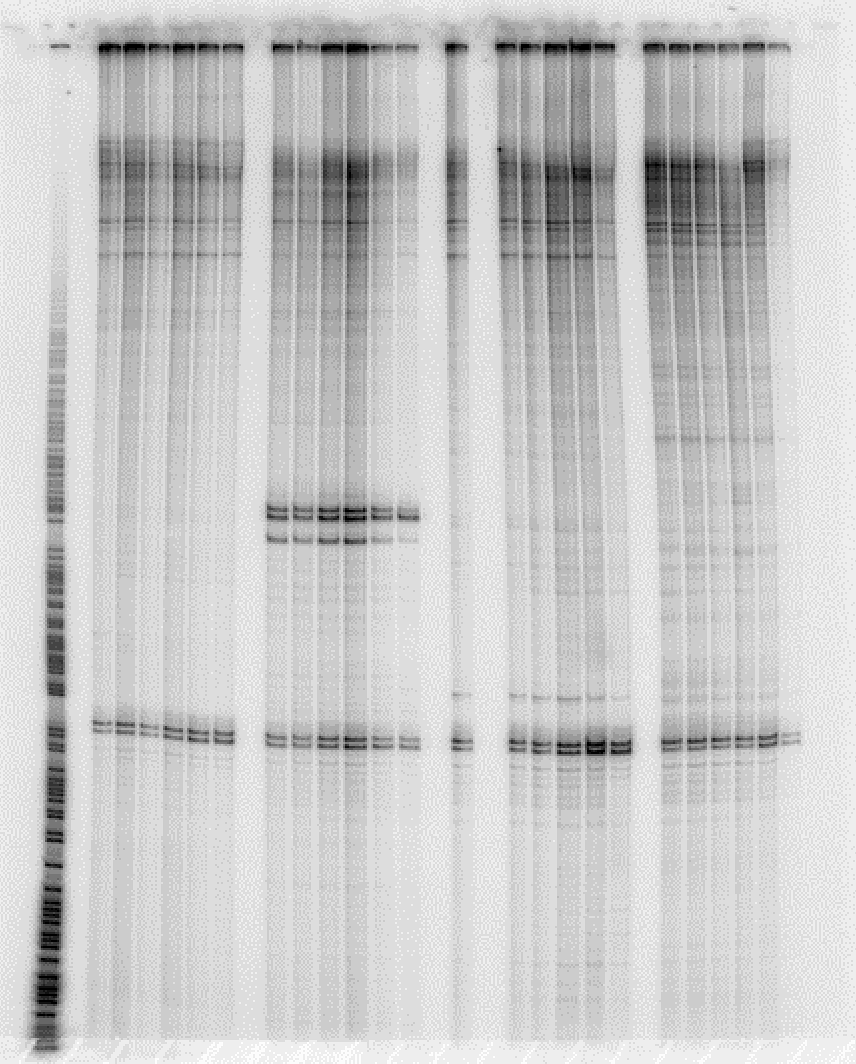

### Figure 2b source data 5

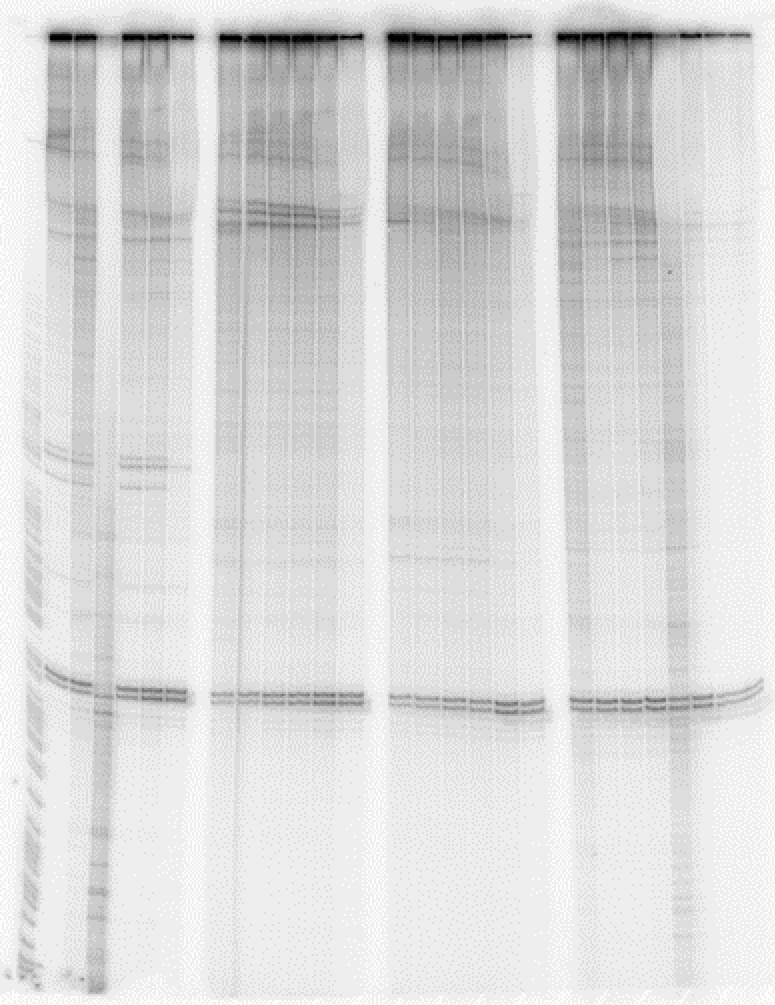

### Figure 2b source data 6

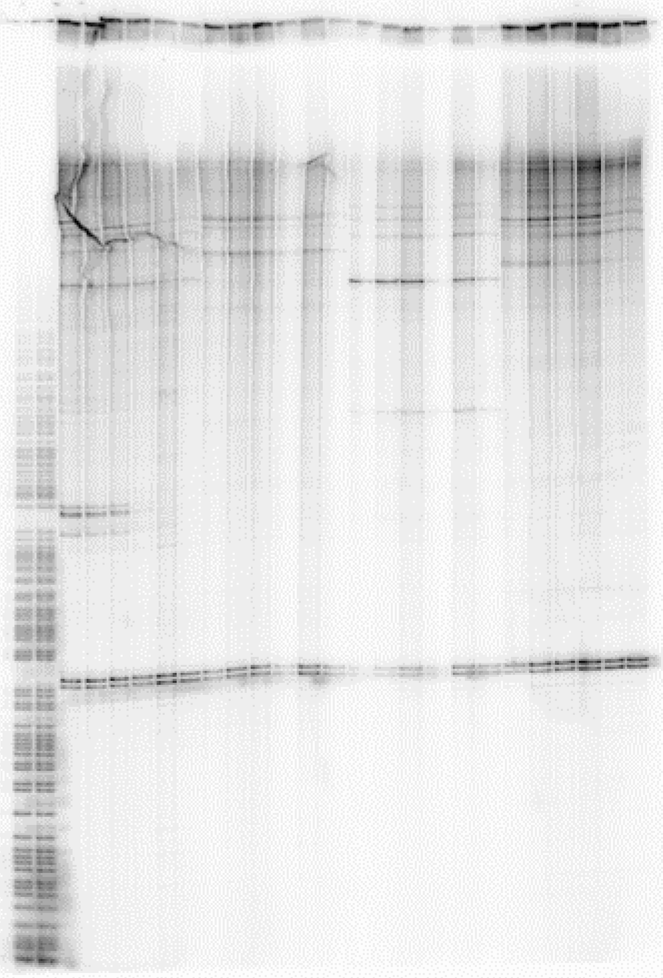

### Figure 3c source data 1

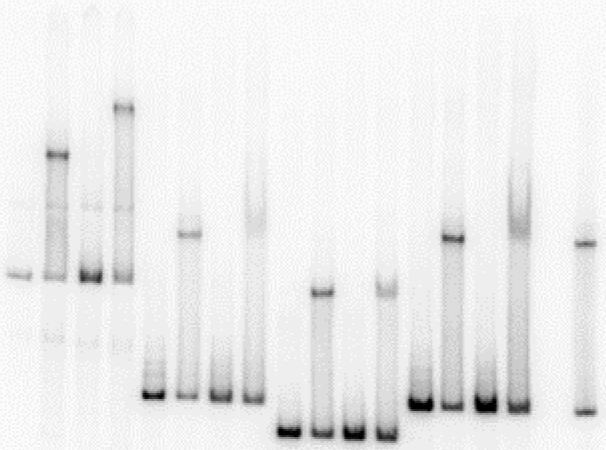

### Figure 3c source data 2

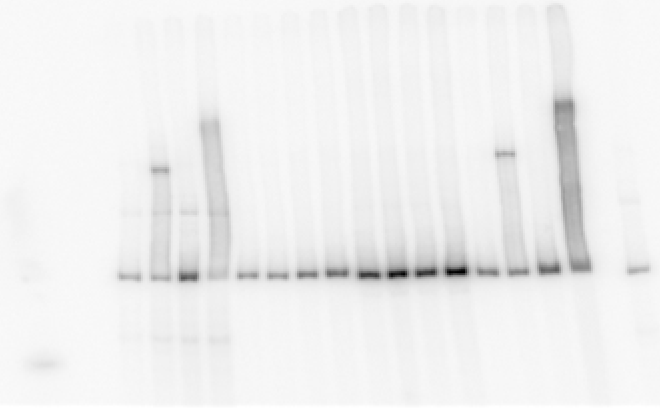

### Figure 3d source data

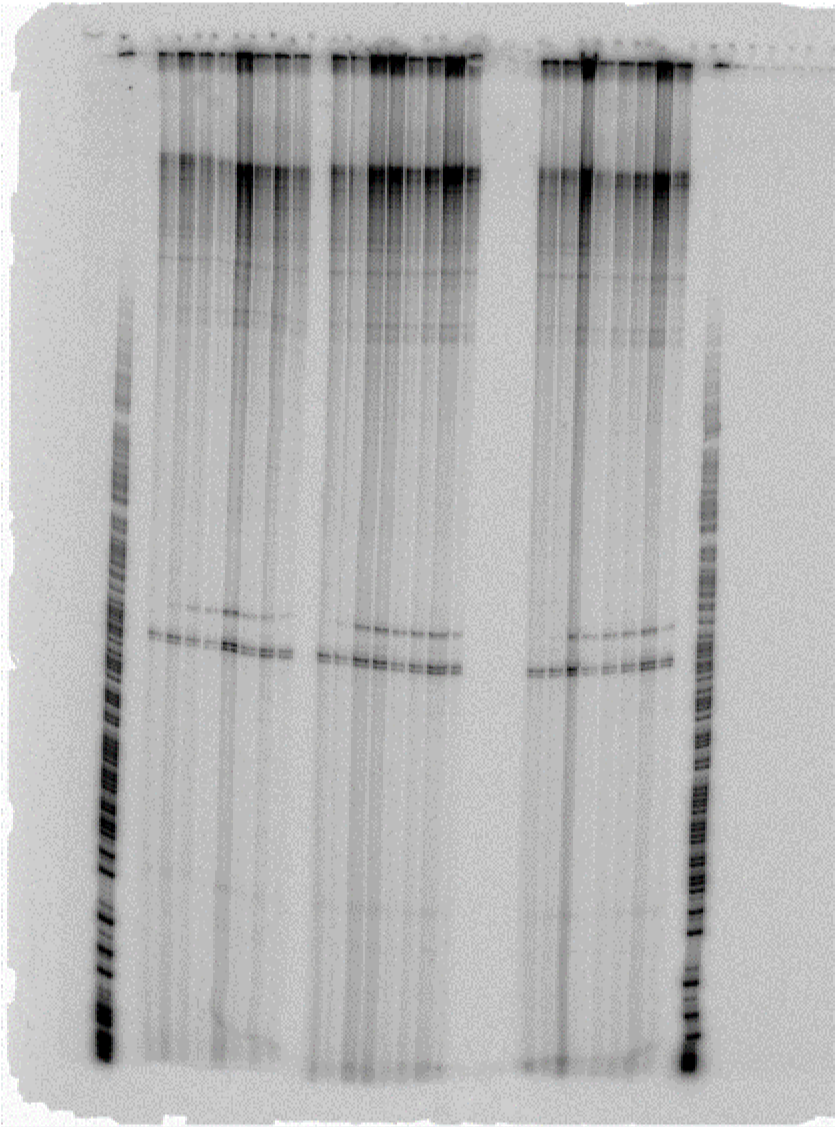

### Figure 4-figure supplement 1

#### Figure 4 figure supplement 1

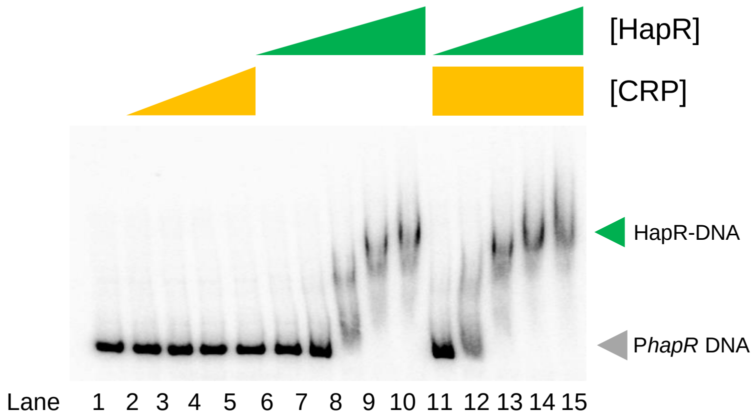

### Figure 4a source data

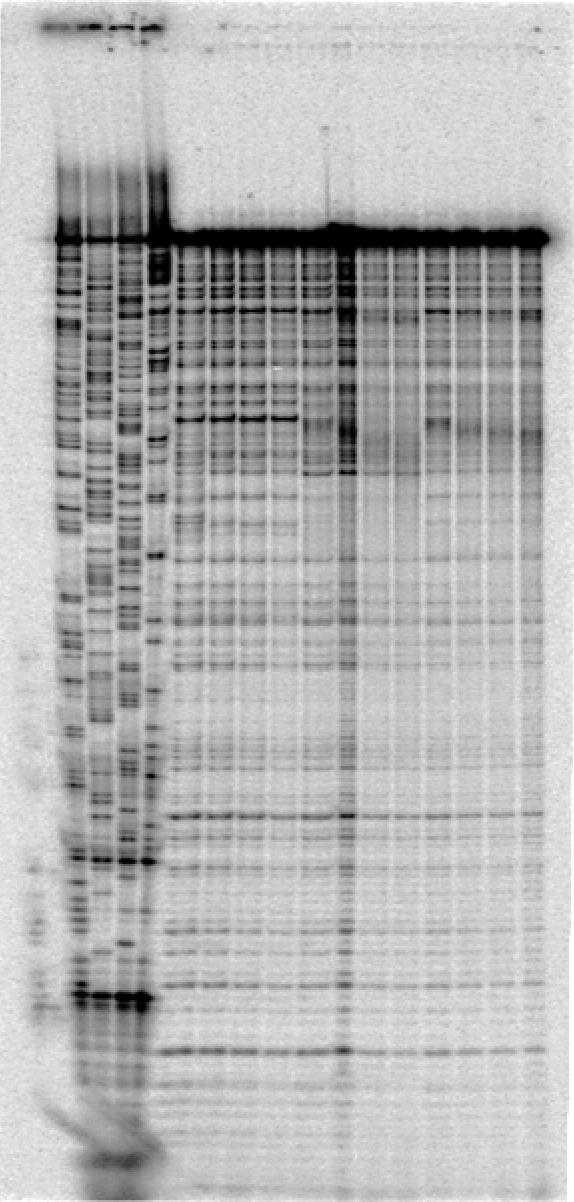

### Figure 4b source data

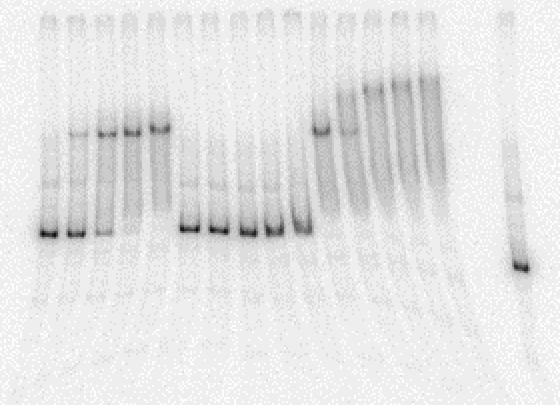

### Figure 4c source data 1

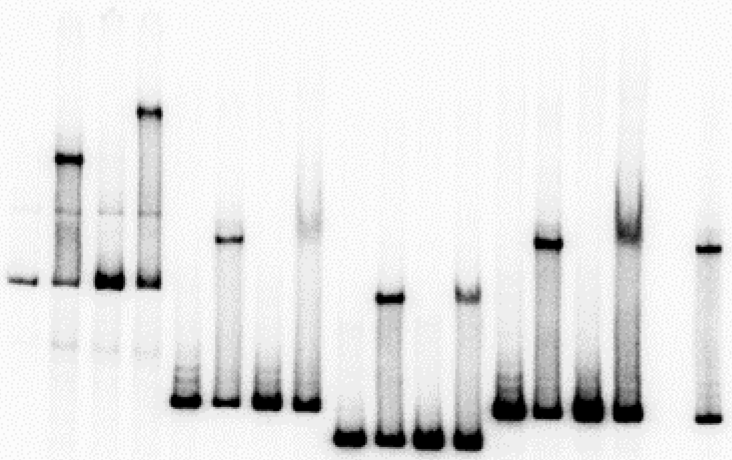

### Figure 4c source data 2

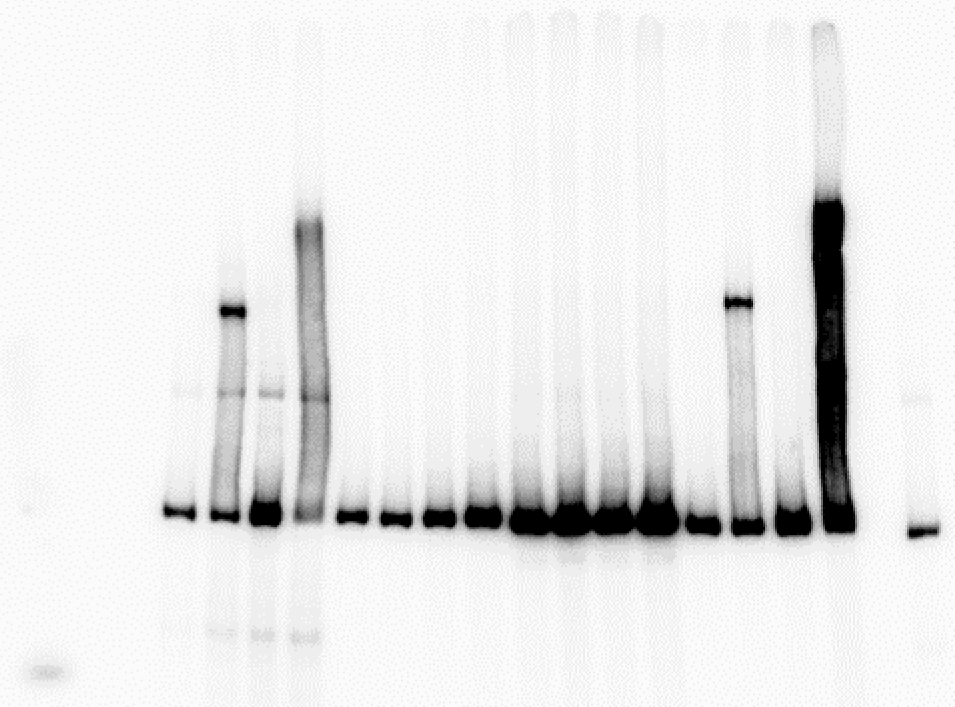

### Figure 4d source data

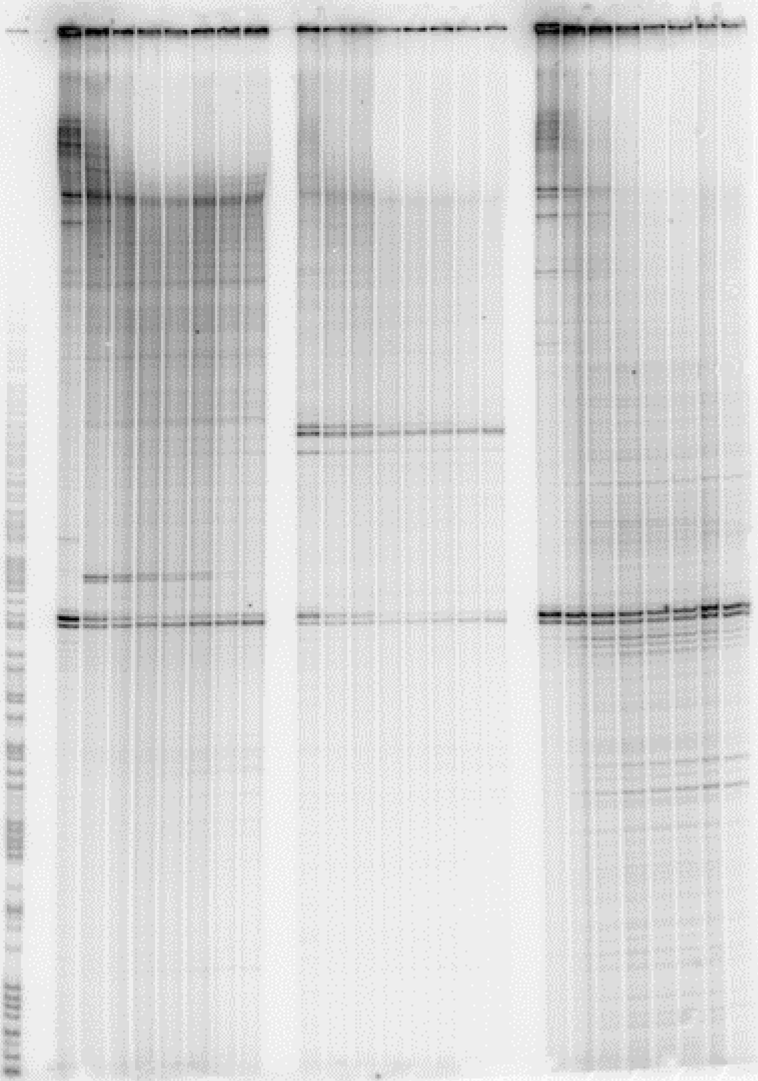

### Figure 5-figure supplement 1

#### Figure 5-figure supplement 1

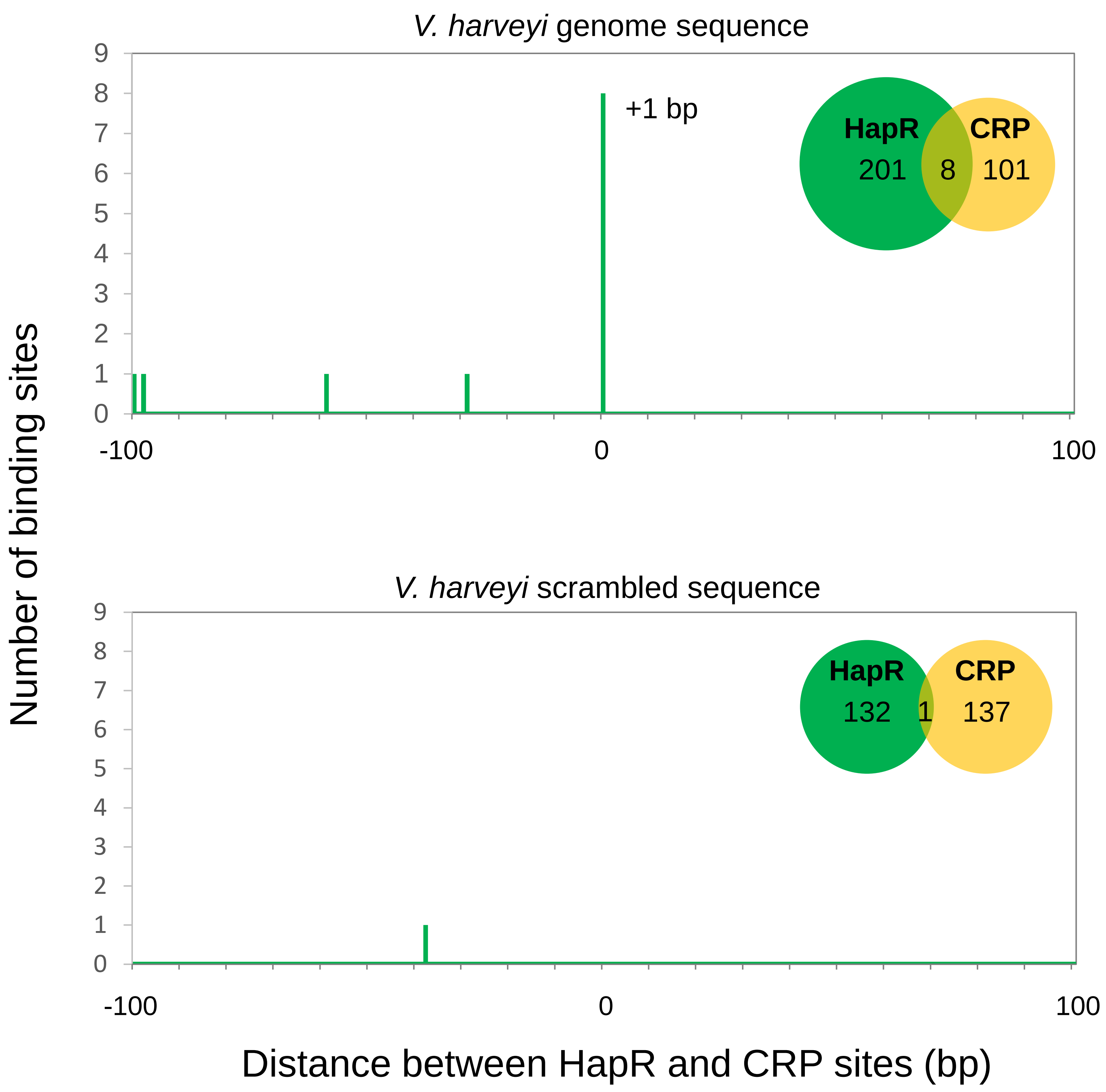

### Figure 5-figure supplement 2

#### Figure 5-figure supplement 2

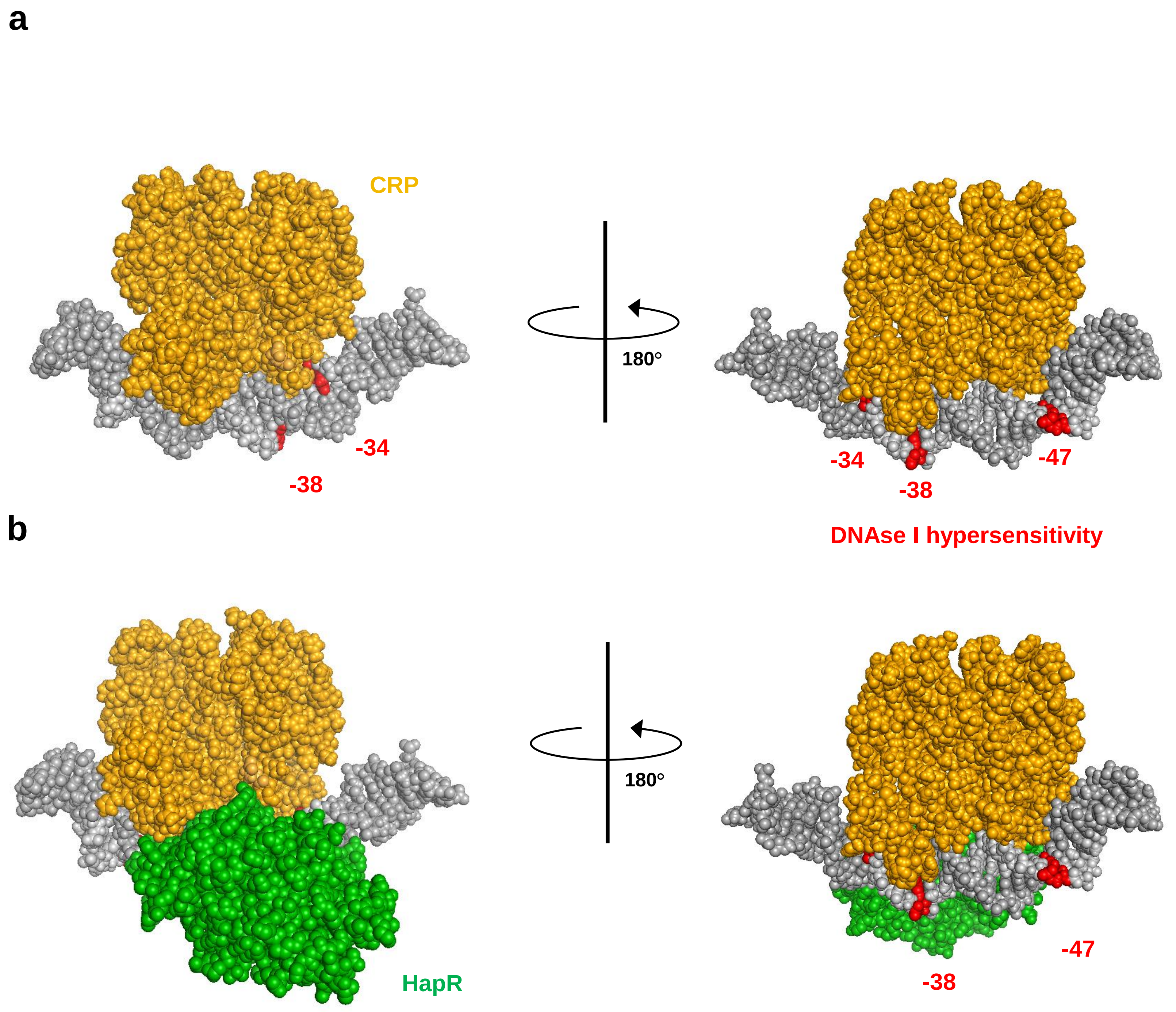

### Figure 5-figure supplement 3

#### Figure 5-figure supplement 3

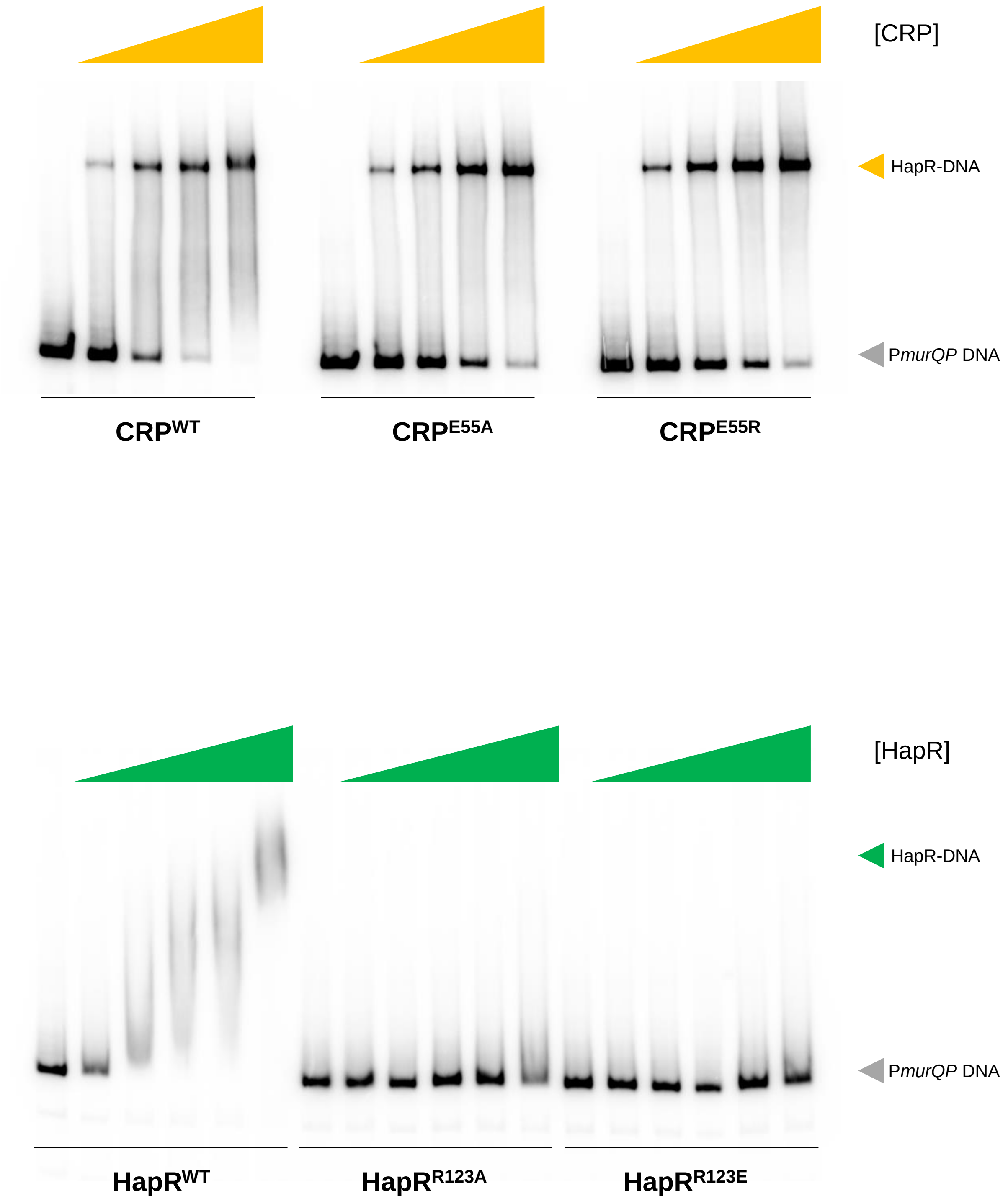

### Figure 5-figure supplement 3 source data 1

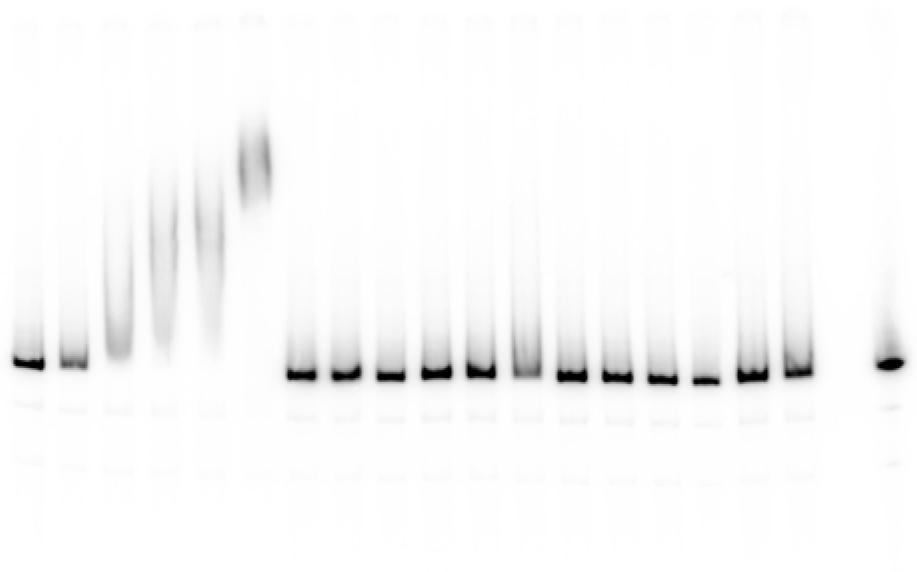

### Figure 5-figure supplement 3 source data 2

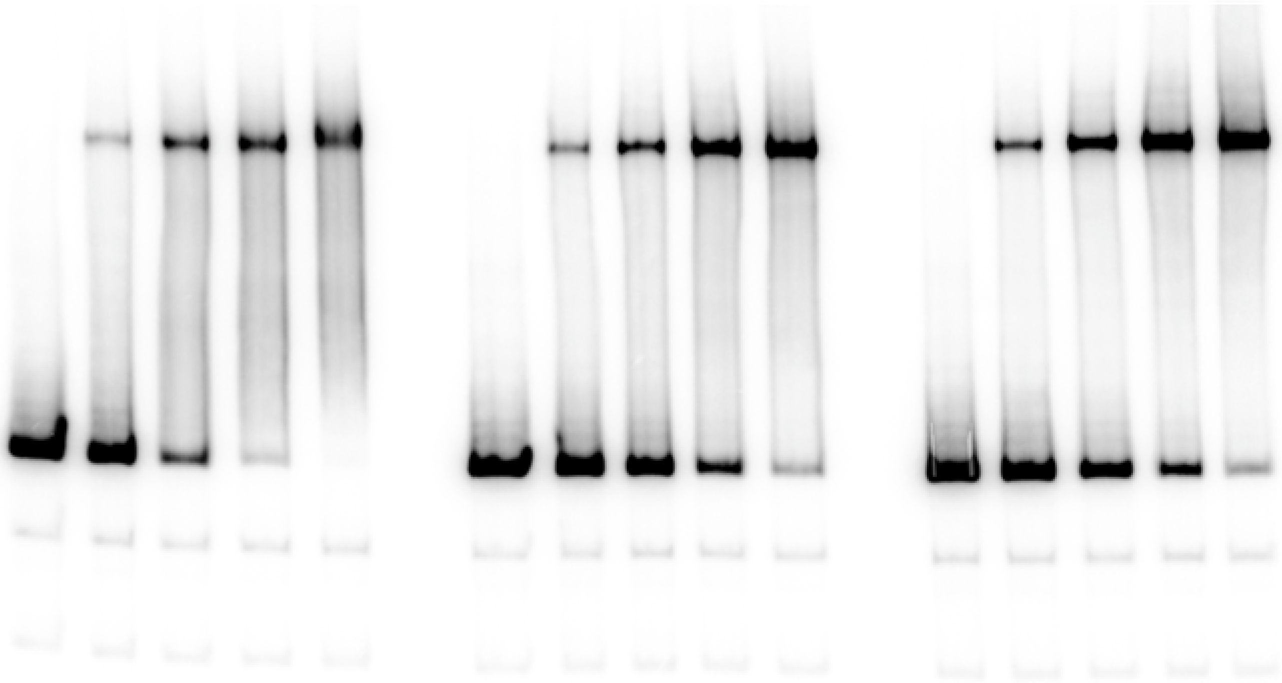

### Figure 5-figure supplement 4

#### Figure 5-figure supplement 4

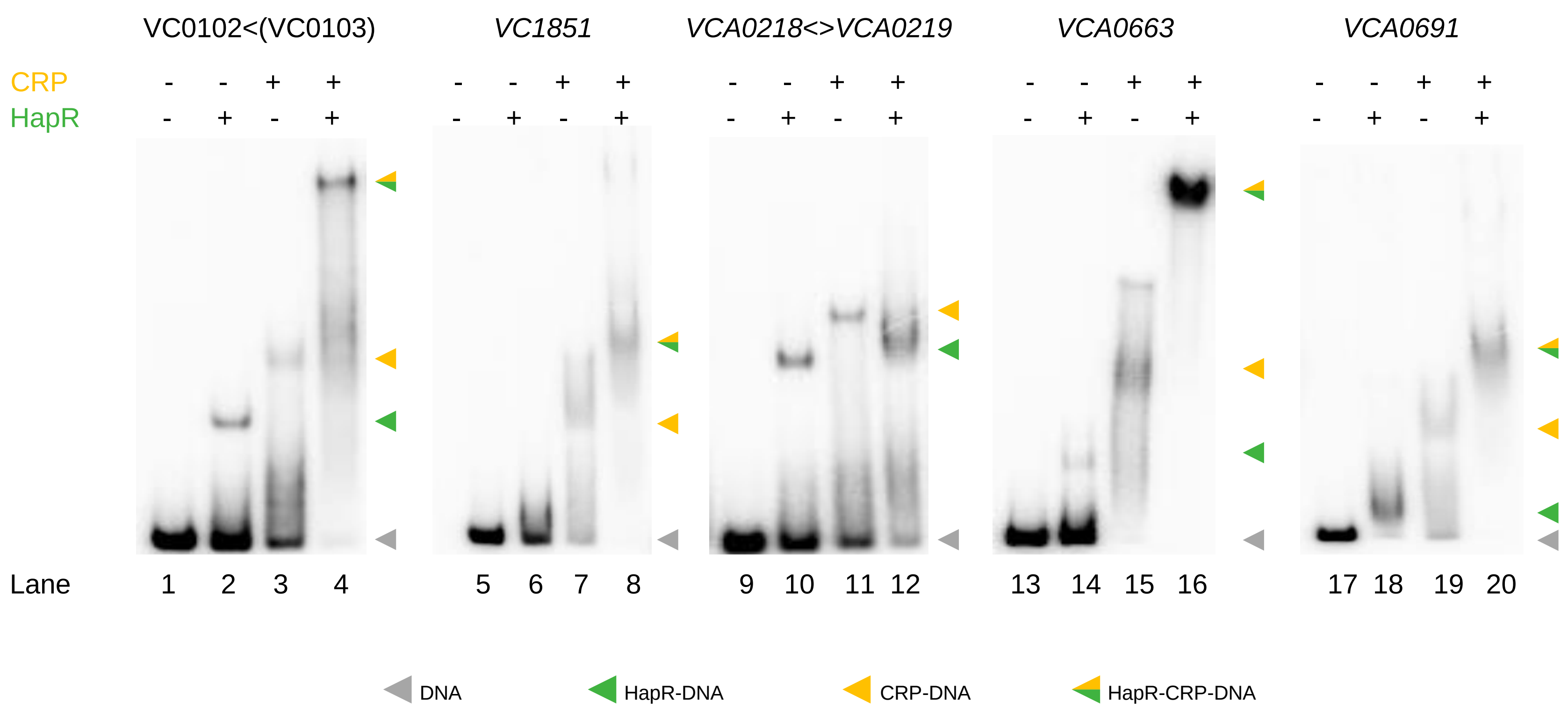

### Figure 5-figure supplement 4 source data 1

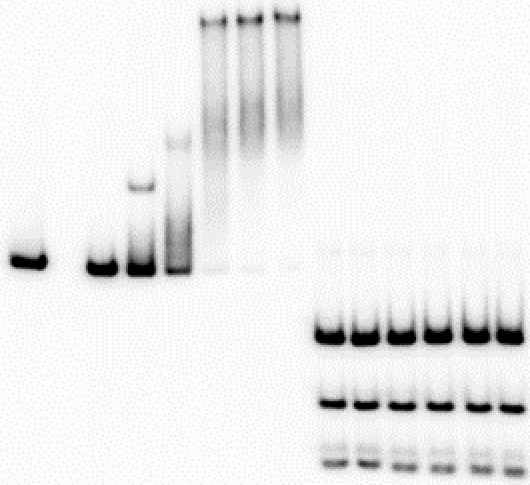

### Figure 5-figure supplement 4 source data 2

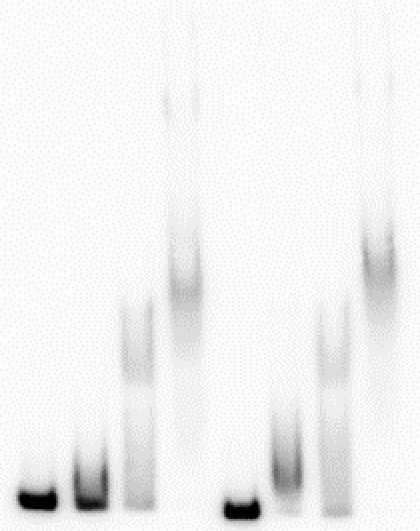

### Figure 5-figure supplement 4 source data 3

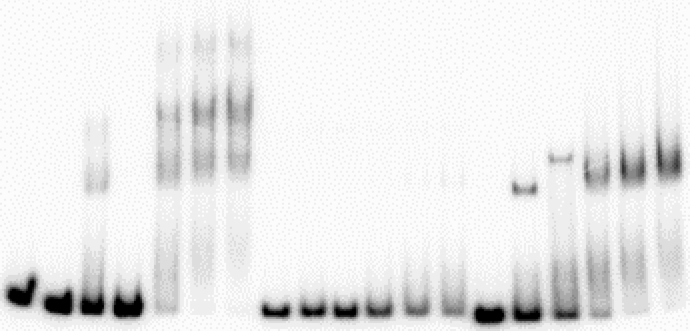

### Figure 5-figure supplement 4 source data 4

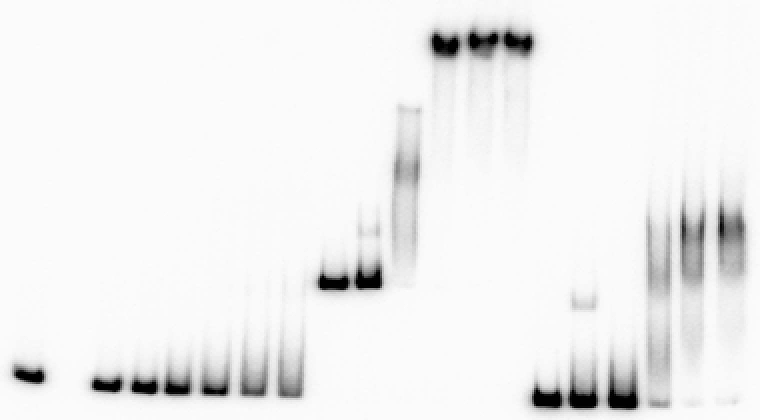

### Figure 5c source data

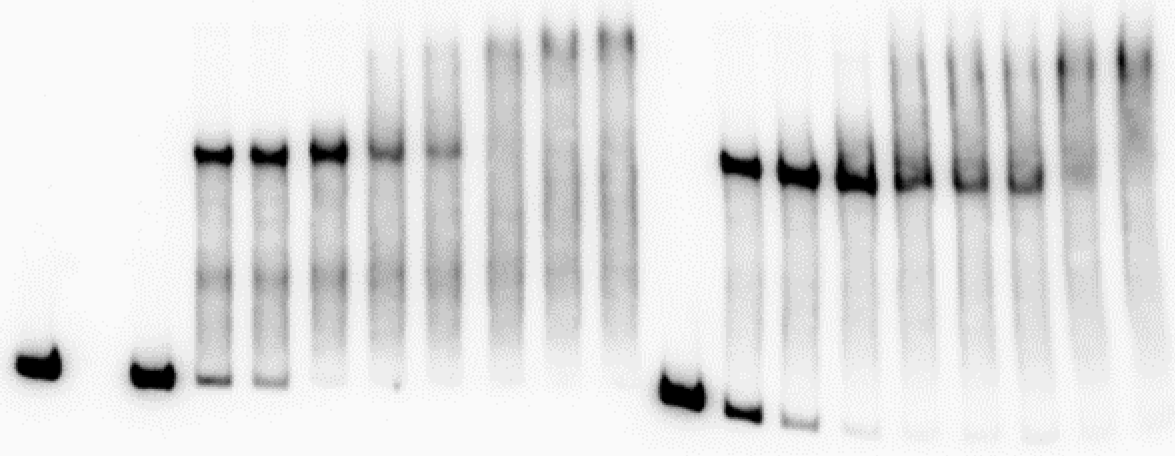

### Figure 5e source data

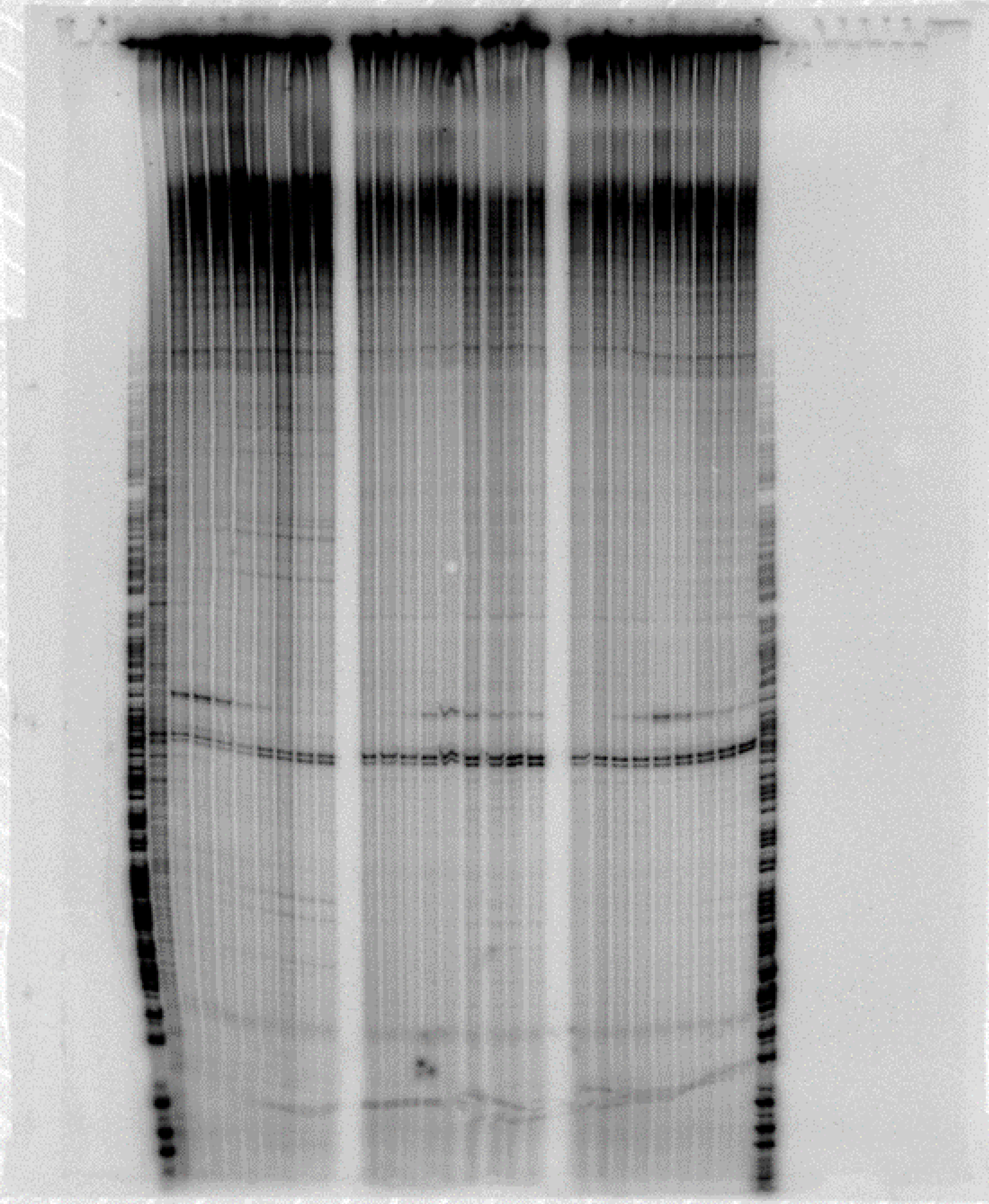
